# siRNA complexation with Galactose-Functionalized Dendrimer: PAMAM vs PETIM

**DOI:** 10.1101/2025.01.18.633705

**Authors:** Tarun Maity, Yogendra Kumar, Prabal K. Maiti

## Abstract

siRNA-based therapies hold promise for treating various diseases, but efficient and targeted delivery remains challenging. This study explored the potential of galactose-functionalized poly(amidoamine) (Gal-PAMAM) and poly(propyl ether amine) (Gal-PETIM) dendrimers as siRNA carriers using all-atom molecular dynamics simulations. This study revealed distinct complexation behavior for both the system in physiological (pH ∼ 7) and high (pH ∼ 10) environments. Electrostatic interactions dominated complexation, with PMF profiles revealing stronger binding affinity for Gal-PAMAM/siRNA at pH 7. siRNA displayed lower flexibility at pH 7 for both complexes, indicating enhanced stability. These findings suggest that Gal-PAMAM dendrimers at physiological pH hold promise for efficient siRNA delivery due to their superior complex stability and interaction strength. However, in polar solvents, Gal-PETIM exhibits hydrophobic behavior, enabling rapid delivery.

## Introduction

Nucleic acid therapies hold significant therapeutic promise for treating a wide range of diseases, including fibrosis,^1^ diabetes,^2^ heart disease,^3^ and cancer, ^4^,^5^ infectious diseases, autoimmune diseases, and HIV/AIDS.^6^ Nucleic acid therapy employs nucleic acids as active agents and can modify gene expression or generate therapeutic proteins, making them well-suited for pathologies with established genetic targets for these diseases.^7^ In recent years, regulatory agencies have approved multiple gene therapy products for various applications, with a notable example being the authorization of mRNA vaccines for COVID-19.^8^

However, despite considerable progress in this field, the transport of large, fragile, and negatively charged molecules, such as short interfering RNA (siRNA), microRNA (miRNA), DNA, and proteins, to their respective targets remains a significant challenge. The inherent challenges associated with viral gene delivery systems have prompted the development of alternative non-viral gene delivery strategies. These nonviral systems typically involve the compaction of negatively charged RNA/DNA molecules into dense, positively charged (or neutral) particles that target cells can effectively internalize via endocytosis. Various nonviral delivery carriers, such as dendrimers,^9–13^ cationic lipids, ^14–16^ carbon nanotubes,^17–21^ proteins,^22–24^ and cell-penetrating peptide, ^25–27^ have been employed to achieve siRNA compaction.

Among these carriers, poly(amido amine) (PAMAM),^12^,^28^ and poly(propyl ether imine) (PETIM)^13,29^ dendrimers stand out due to their well-defined three-dimensional structure and a multivalent terminal group that can be easily tuned.^28,30^ This dendritic architecture and multivalent polycationic surface group render dendrimers highly efficient drug delivery vehicles.^31,32^ Experimental and simulation studies suggest that complexes formed between siRNA and dendrimers^33–42^ and between DNA and PAMAM dendrimers^43–47^ exhibit higher transfection efficiency than other conventional gene delivery systems. The functionalization of dendrimers, particularly PAMAM and PETIM, further enhances their potential as gene delivery agents.^48,49^ By attaching peptides or small biological molecules to their surface groups, researchers can tailor these carriers for targeted delivery to specific cells or tissues, including cancer cells. ^29^

In the context of cancer treatment, targeted delivery is essential.^50^ Due to their rapid metabolism, cancer cells consume more glucose than normal cells to sustain their continuous growth.^51,52^ This increased glucose uptake, is facilitated by the over expression of GLUT receptors on cancer cell surfaces. These findings present an opportunity for targeted nanoparticle drug delivery systems. Indeed, glucose-modified nanoparticles have shown significantly enhanced cellular uptake compared to unmodified ones.^53^

Recently, sugar-functionalized dendrimers have emerged as promising agents for targeting cells.^54–57^ The glycosylation of the PAMAM dendrimer demonstrated that glucose modification facilitated the targeting of microglia and tumor-associated macrophages (TAMs) by augmenting cellular internalization brain penetration. ^57^ The galactose-modified PAMAM dendrimer is more efficient in targeting the glioblastoma tumor cells (brain cancer) than glucose and mannose modifications.^57^ The n-acetylgalactosamine ligand was attached to the surface group of the PAMAM dendrimer, which successfully delivered the siRNA to the target cell and showed reduced toxicity. ^58^ These findings highlight the versatility and efficacy of sugar-functionalized dendrimers for targeted drug and gene delivery applications. However, a detailed understanding of the interactions between siRNA and galactose functionalized dendrimer complex at the molecular level is lacking, hindering further optimization of these delivery systems. Recent studies with unmodified PAMAM dendrimer highlighted the impact of pH on siRNA complex formation, but the influence of galactose modification on these interactions remains unexplored. Here, we reveal molecular details of the siRNA binding to the galactose-functionalized dendrimer complex. ^57^

We organize the rest of the paper as follows: Section provides details of the simulation models and methodology used in this work. Section presents key results on intermolecular interactions between Gal-dendrimer and siRNA at different pH values. We also discuss the structural properties and kinetics of complex formation. Section summarizes the key findings on intermolecular interactions and complex formation between Gal-dendrimer and siRNA. It also provides insights into potential future research directions.

## Materials and methods

### Model Building

Galactose-functionalized PETIM (Gal-PETIM) and PAMAM (Gal-PAMAM) dendrimers were constructed using the following approach. Dendrimers comprise three fragments: core, repeating units (monomers), and surface groups. The third-generation (G3) Gal-PETIM and Gal-PAMAM dendrimers were constructed directly from the core, monomer, and galactose-functionalized surface group using our in-house dendrimer building toolkit (DBT).^59^ The galactose molecule was geometrically optimized using the Gaussian09 software package^60^ with Hartree-Fock (HF) level of theory and 6-31G(d) basis set. Partial atomic charges were assigned using the restrained electrostatic potential (RESP) method as implemented in the antechamber module of AMBER20.^61^ The galactose molecules were covalently attached to the surface amine groups of dendrimer using ChemDraw 22.2.0 software. Partial atomistic charges of the core and monomer units were adopted from our previous work.^13,59,62^ The initial configuration for the 21-base pair siRNA was obtained from the Protein Data Bank (PDB ID: 2F8S).^13,63^ The AMBER14SB force field with PARMBSC1^64^ corrections were used to describe the interactions involving the siRNA molecule. The general amber force field (GAFF)^65^ force field parameters were employed to describe the inter- and intramolecular interaction involved in Gal-PAMAM/Gal-PETIM dendrimers.

All the atomistic MD simulations presented in this work were carried out using GROMACS (version 2022.5) package. ^66,67^ The galactose-functionalized dendrimers (Fig. 1) along with siRNA was placed in a 10 *×* 10 *×* 10 nm^3^ periodic cubic box (Fig. 2). Subsequently, solute (siRNA and Gal-PETIM/Gal-PAMAM) was solvated using the TIP3P water^68^ model. Appropriate numbers of Cl^−^ and Na^+^ counterions were added to neutralize the positive charges on protonated Gal-PETIM/Gal-PAMAM (G3) dendrimers and negative charges on the phosphate backbone of the siRNA, respectively. The initial solvated and charge-neutralized Gal-PAMAM/siRNA system is shown in Fig. 2. The solvated systems were then subjected to energy minimization using 1000 steps of the steepest descent algorithm to remove high-energy interactions from the initial configuration. Subsequently, the systems were heated to 303 K using Noose-Hoover thermostat^69,70^ in constant temperature-constant volume (NVT) ensemble with a damping time constant 1.0 ps. During heating, the solute was kept fixed at its minimum energy configuration, using harmonic restraint with force constant 1000 kJmol^*−*1^nm^*−*2^. After heating, the equilibrium MD simulations were performed at NPT ensemble for 1 ns to achieve the equilibrated density using the Parrinello-Rahman^71^,^72^barostat with relaxation time of 5.0 ps. We then performed an MD production run under constant pressure (p = 1 bar) and temperature (T = 303 K) conditions (NPT) for 200 ns. The pressure was achieved by applying Parrinello-Rahman^71^,^72^ barostat with a relaxation time of 5.0 ps. Temperature regulation was achieved Nosé–Hoover thermostat with a damping time constant 1.0 ps. We considered water compressibility of 4.5*×* 10^*−*10^ kPa^*−*1^. LINCS^73^ algorithm was used to constrain all covalent bonds involving hydrogen atoms. Long-range electrostatic interaction was calculated using particle mesh Ewald (PME) method^74^,^75^ with a cut-off 12 Å. The short-range van der Waal interactions were calculated using the Lennard-Jones (LJ) potential with a cutoff of 12 Å. Visual molecular dynamics (VMD)^76^ was used to visualize the trajectories and render the snapshots.

**Figure 1.**
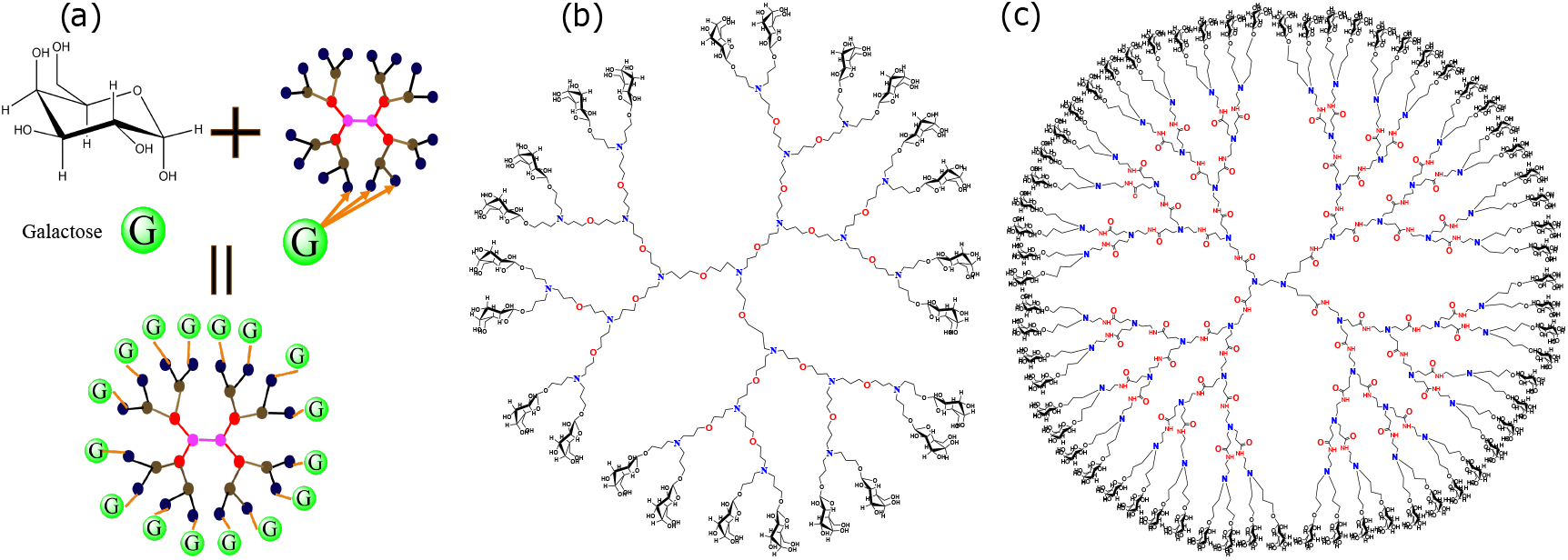
Chemical structure of galactose and a schematic representation of its conjugation with the dendrimer.’G’ represents galactose molecules. Chemical structures of (b) galactose functionalized third-generation (G3) nitrogen-cored PETIM and (b) galactose functionalized third-generation ethyl diamine-cored PAMAM dendrimer.

**Figure 2.**
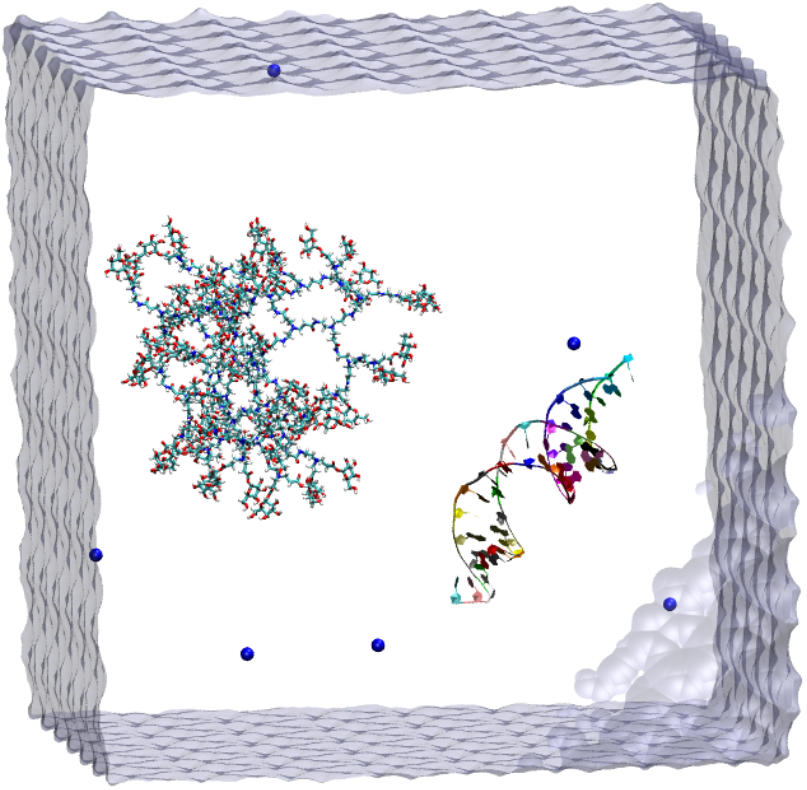
Initial solvated system of Gal-PAMAM/siRNA at pH 7, with a center-of-mass separation (COM-COM) distance of 5 nm between the dendrimer and siRNA. Color code: water in ice-blue, Na^+^ in blue vdW ball, Gal-PAMAM(G3) in Licorice, and siRNA in NewRibbons with resid representation

### Umbrella Sampling Simulations

To investigate the free energy landscape of siRNA-Gal-dendrimer dissociation, we employed the umbrella sampling (US) technique to calculate the potential of the mean force (PMF) profile along the dissociation pathway. PMF quantifies the system’s free energy as a function of a chosen reaction co-ordinate (RC)(*ξ*). RC was defined as the center-of-mass (COM) distance between the Gal-PAMAMA/Gak-PETIM and siRNA. Using steered molecular dynamics (SMD), Gal-PETIM/Gal-PAMAM was pulled to dissociate by moving its COM along the RC to generate the initial configuration for the US simulation. The pulling rate of the Gal-PETIM/Gal-PAMAM molecules was 0.01 nm/ps with a spring constant of 1000 kJmol^*−*1^nm^−2^. We have sampled the RCs starting from the equilibrium distance in the complex state and extending up to 60 Å in steps of 1 Å to capture the dissociation process fully. From the SMD run, we have generated 150− 212 windows for umbrella sampling. Each window was simulated for 5 ns with a harmonic bias potential of the force constant of 1000 kJmol^−1^nm^−2^ to obtain a proper probability distribution for each window and the convergence of the PMF profile. After obtaining the histograms corresponding to each window of the umbrella sampling simulation, the weighted histogram analysis method (WHAM)^77^ was used to obtain the PMF profiles.

## Results and Discussion

We observe the complex formation of Gal-PAMAM/Gal-PETIM and siRNA at both pH values (pH 7 and pH 10), as can be seen from the instantaneous snapshots shown in Fig. 3 (a-h). Intriguingly, a notable difference in complex morphology was observed between the two pH environments. While complexes formed in all simulations, Fig. 3 (a-h)) reveals a more extensive coverage of the siRNA surface by the Gal-PETIM/Gal-PAMAM dendrimer at pH 7 (Fig. 3 (a-d))) compared to pH 10 Fig. 3 (e-h)). This observation suggests stronger intermolecular interactions between Gal-PETIM/Gal-PAMAM and siRNA at physiological pH (pH ∼ 7), which may have implications for the efficiency of siRNA delivery in biological systems.

**Figure 3.**
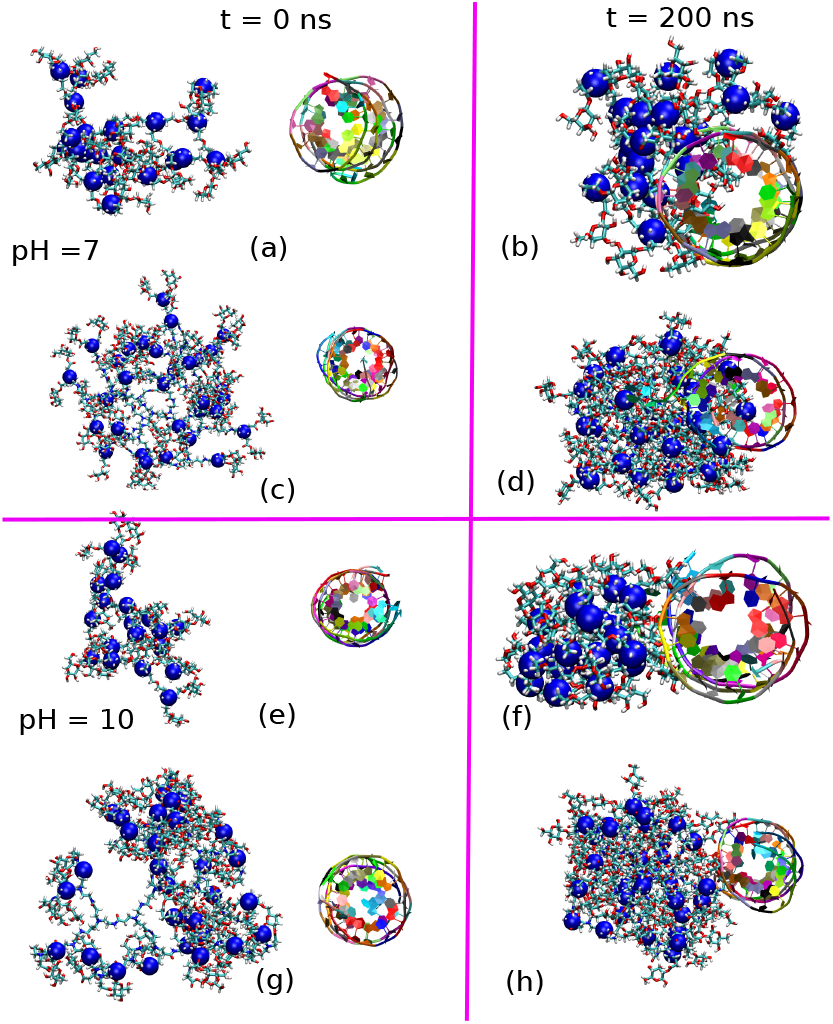
Snapshots of (a,b,e,f) the Gal-PETIM(G3)/siRNA and (c,d,g,h) the Gal-PAMAM(G3/siRNA at pH 7 and pH 10, respectively. Color code: blue beads are surface amine nitrogen, and the rest of the dendrimer is in Licorice VMD representation, siRNA is in NewRibbons with resid representation

### Center-of-mass (COM) distance between siRNA and the Gal-PAMAM/Gal-PETIM

At pH 7, protonated surface amines on the dendrimer electrostatically attract the negatively charged phosphates group of siRNA, facilitating rapid complexation. In contrast, deprotonation of these amines at pH 10 results in weaker, predominantly van der Waals interactions, leading to delayed and weaker complexation.

Fig. 4 depicts the time evolution of the center-of-mass (COM) distance between siRNA and (a) Gal-PETIM (b) and Gal-PAMAM at two different pH conditions (pH 7 and 10). A rapid decrease in COM distance is observed at pH 7 for both dendrimers, indicating swift complexation between the dendrimer and siRNA. The subsequent plateauing of the COM distance suggests stable complex formation. For Gal-PETIM(G3)-siRNA exhibited a rapid decrease in the COM distance from 5 nm to 1.2 nm within 20 ns, indicating fast complexation, followed by a plateau, which is indicative of complex stability (Fig. 4 (a)). Gal-PAMAM (G3) formed complex within 15 ns settling at a distance of 1.4 nm. In contrast, at pH 10, complexation is slower for both dendrimers. For Gal-PETIM(G3)-siRNA, the COM distance between siRNA and Gal-PETIM fluctuated until 30 ns before decreasing to 2.2 nm by 80 ns. Gal-PAMAM (G3) formed a stable complex within 40 ns, and the COM distance fluctuates around 4 nm. This implies that the interaction between the dendrimers and siRNA is pH-dependent, with stronger and more stable complexation occurring at physiological pH (= 7). In contrast, deprotonation of these amines at pH 10 results in weaker, predominantly van der Waals interactions, leading to delayed complexation and larger fluctuations in COM distance.

**Figure 4.**
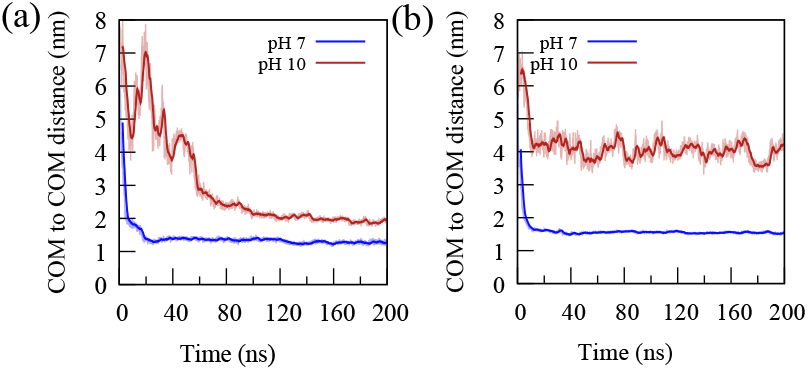
Temporal change in center-of-mass distance between the (a) Gal-PETIM/siRNA and (b) Gal-PAMAM/siRNA complex at two different pH values.

### Number of Contacts and Hydrogen Bonding

To investigate the influence of pH on the binding affinity between Gal-PAMAM/Gal-PETIM and siRNA, we computed the time evolution of the number of close contacts (*N*_*c*_) - defined as the number of dendrimer atoms within 0.3 nm of any siRNA atom. Consistent with the rapid decrease in the center-of-mass (COM) distance at pH 7 (Fig. 4), a significant increase in *N*_*c*_ was observed for both Gal-PETIM and Gal-PAMAM dendrimers (Fig. 5a, b). This reflects closer proximity and stronger interactions, primarily attributed to the electrostatic attraction between positively charged dendrimer surface groups and negatively charged siRNA phosphate groups. In contrast, at pH 10, *N*_*c*_ remained lower due to the deprotonation of the amine groups, resulting in weaker van der Waals interactions.

**Figure 5.**
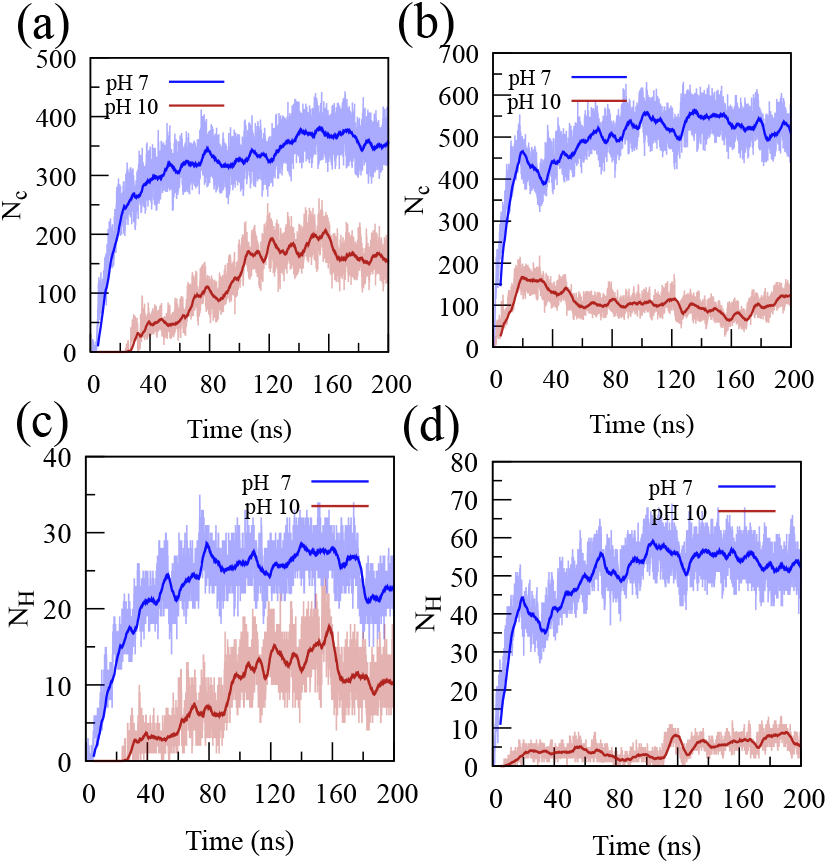
Temporal variation in (a,b) the number of contacts (*N*_*c*_) and (c,d) the number of hydrogen bonds (*N*_*H*_) at two different pH values (pH 7 and 10) between the siRNA and galactose functionalized PETIM and PAMAM dendrimers.

We have also computed the number of hydrogen bonds (*N*_*H*_) between the siRNA and dendrimer. The number of hydrogen bonds (*N*_*H*_) between the dendrimers and siRNA was calculated using a geometric criterion with a distance cutoff of 0.35 nm and an angle cutoff of 120^°^. Notably, Gal-PAMAM exhibited a higher *N*_*H*_ than Gal-PETIM, suggesting stronger binding affinity at physiological pH. This is due to the larger number of available binding sites at G3 PAMAM compared to the G3 PETIM. At pH 10, *N*_*H*_ for Gal-PETIM initially increased up to 150 ns before stabilizing, while Gal-PAMAM showed a lower and less stable complex.

### Conformational Dynamics of siRNA-Gal-Dendrimer Complexes

The efficiency of siRNA delivery depends not only on binding affinity but crucially on the size of the resulting complexes. To quantify the size of the complex, we have calculated the radius of gyration (*R*_*g*_) of both the isolated dendrimers and the siRNA-dendrimer complexes at different pH values (Fig. 6). The *R*_*g*_ values reflect how the dendrimers adjust their conformation upon complexation with siRNA. Figure 6 reveals fluctuations in *R*_*g*_ during the initial complexation phase (up to 20 ns), likely due to the dendrimers optimizing electrostatic interactions with siRNA. After complex formation, *R*_*g*_ stabilizes for Gal-PETIM(G3) and Gal-PAMAM(G3). Figure 6 depicts that Gal-PAMAM dendrimers are larger than their Gal-PETIM at the same generation under both pH conditions. The size difference is attributed to the greater number of terminal groups in the PAMAM dendrimer (2^(*G*+2)^ scaling) compared to the PETIM dendrimer (2^(*G*+1)^ scaling). ^78^ For G3 dendrimers, this translates to 32 terminal groups for PAMAM, leading to a more open and expanded structure compared to PETIM (12 terminal groups).

**Figure 6.**
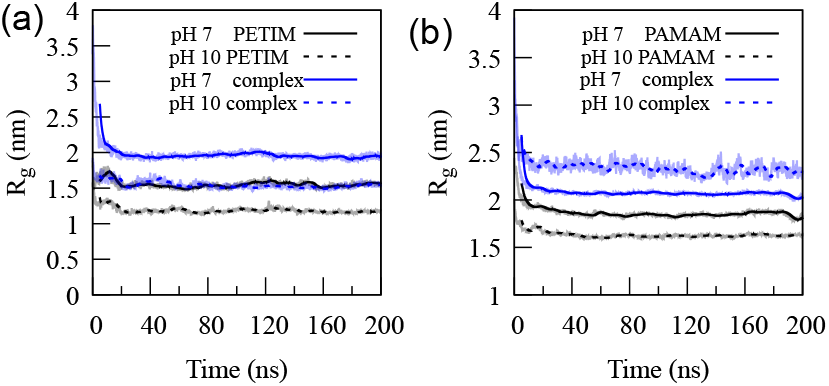
Time series plot of radius-of-gyration (*R*_*g*_) of (a) Gal-PETIM and Gal-PETIM/siRNA complex, and (b) Gal-PAMAM and Gal-PAMAM/siRNA complex at the two different pH conditions.

The average *R*_*g*_ values are calculated over the final 100 ns of the trajectory. At physiological pH (= 7), the size of Gal-PETIM (G3) is larger (*R*_*g*_ = 1.56 *±* 0.03 nm) than the size at pH 10 (*R*_*g*_ = 1.17 *±* 0.024 nm). Similar to Gal-PETIM (G3), Gal-PAMAM (G3) also exhibited a pH-dependent size change. The size of Gal-PAMAM (G3) at pH 7 is larger (*R*_*g*_ = 1.84 *±* 0.061 nm) than the size at pH 10 (*R*_*g*_ = 1.66 *±* 0.014 nm). This is due to the strong electrostatic repulsion between the protonated surface groups at physiological pH.

### Structure and stability of the Gal-PAMAM/Gal-PETIM and siRNA complex

We have also computed root-mean-square deviation (RMSD) of the dendrimers and rootmean-square fluctuation (RMSF) of the siRNA to investigate the conformational stability and flexibility of the complexes under different pH conditions (Fig. 7). The RMSD provides a measure of the average deviation of the positions of dendrimer atoms from their initial positions, reflecting the overall structural stability of the dendrimer. RMSF, on the other hand, quantifies the fluctuations of individual residues of siRNA, revealing regions with higher or lower flexibility. At pH 7, the Gal-PETIM/siRNA complex exhibited minimal fluctuations in RMSD, indicating a stable complex (Fig. 7a). In contrast, at pH 10, the Gal-PETIM/siRNA complex displayed significantly larger fluctuations in dendrimer RMSD, suggesting a less stable complex due to weaker interactions between the siRNA and dendrimer (Fig. 7a).

**Figure 7.**
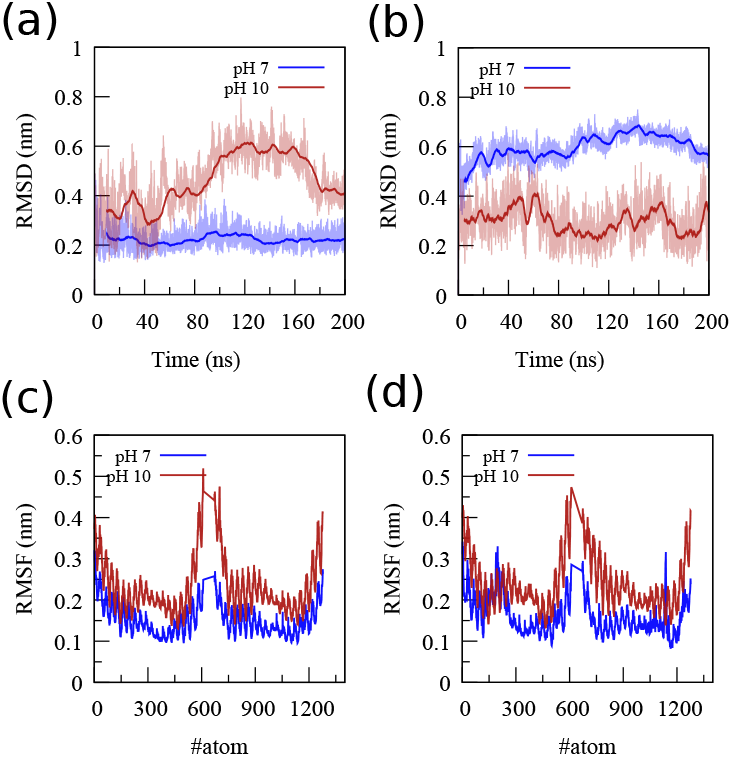
Root mean square deviation (RMSD) and root mean square fluctuation (RMSF) of the siRNA in (a,c) Gal-PETIM/siRNA complex and in (b,d) Gal-PAMAM/siRNA complex at pH 7 and 10, respectively.

The Gal-PAMAM-siRNA complex displayed more structural fluctuations than did the Gal-PETIM-siRNA complex at pH 7. This is attributed to the higher number of charged surface groups on the Gal-PAMAM dendrimer, which led to increased siRNA wrapping and a more pronounced deviation from its initial structure, as evidenced by higher RMSD values. This suggests that complex formation with the dendrimers enhances siRNA stability at physiological pH, likely due to stronger electrostatic interactions.

### Hydrophobic nature: Gal-PAMAM vs Gal-PETIM

Quick delivery of the gene to the targeted cell may be facilitated by the hydrophobic or hydrophilic nature of the dendrimers. We have quantified this by calculating the number of water molecules in different domains of Gal-PAMAM and Gal-PETIM based on the criteria developed earlier in our work. ^79^,^80^Three different domains were defined, namely buried (or internal), surface (or bound), and bulk, for the estimation of the number of water molecules in galactose functionalised dendrimers. The solvent accessible surface area (SASA) of the dendrimers was calculated using the probe radius four times higher than that of water molecules, *i*.*e*. 6 Å to distinguish the internal cavity from the surface. This allows us to define the solvent excluded surface (SES) of the dendrimer. Subsequently, the surface atoms were defined as any atoms that have a nonzero SASA. The surface domain is identified as the space within a spherical radius of 2 Å from any surface atom. Similarly, the domain whih is 3 Å away from the surface atoms is defined as the buried domain. The water molecules within 3 Å of the surface atoms (towards the core of the dendrimers) are defined as buried water. The surface or bound water is calculated 4 Å outside of the surface atoms, and the bulk bound waters are the water molecules that lie between 4 Å and 7 Å from the surface atoms of the dendrimers. Further details on these regimes can be found in Ref.^79^ The results are plotted in Fig. 8 for two different pH conditions.

**Figure 8.**
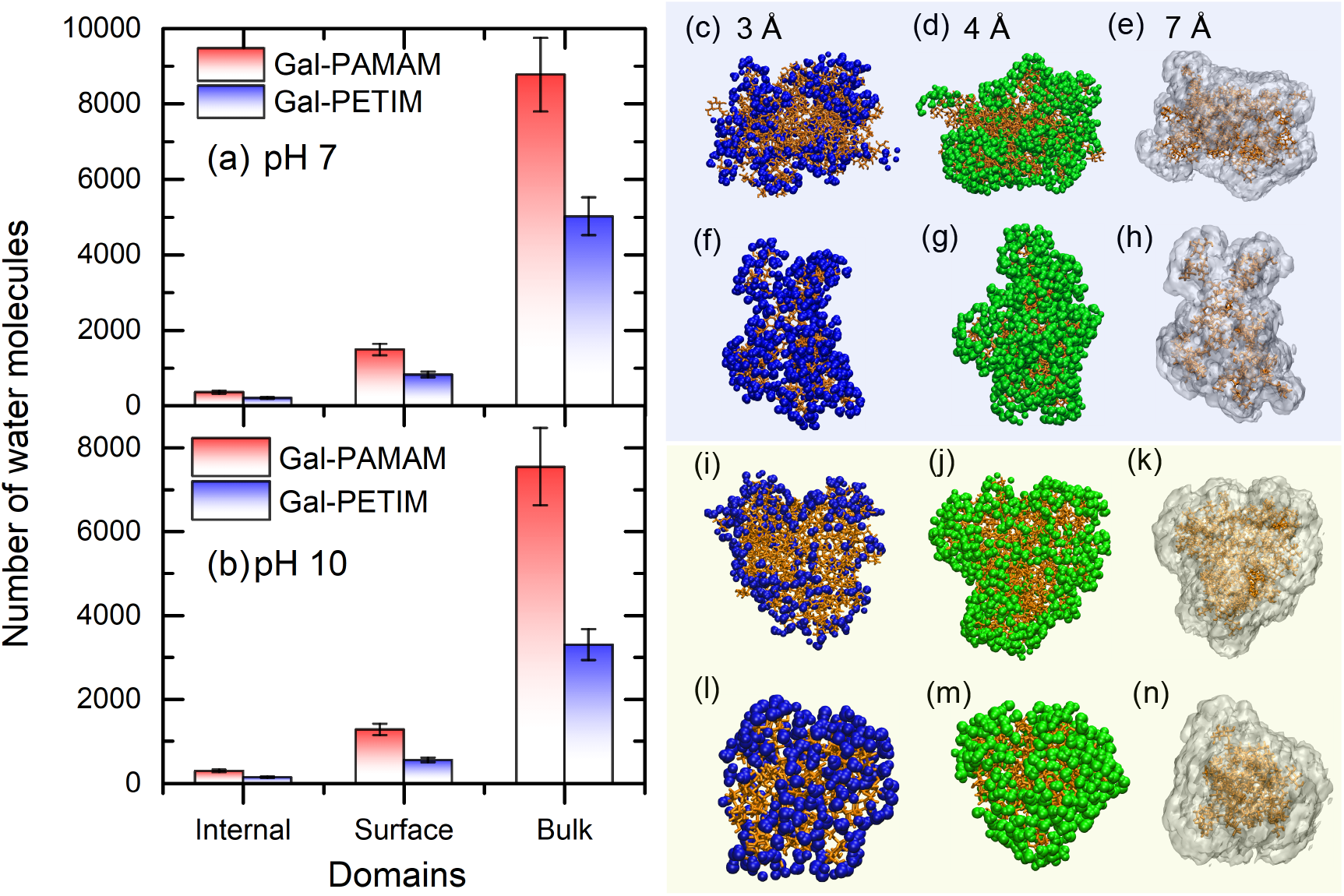
Water at different regimes of Gal-PAMAMA and Gal-PETIM at (a) pH 7 and (b) pH 10. The snapshots in the light blue rectangular box are (c-e) for Gal-PAMAM and (f-h) for Gal-PETIM at pH 7, while in the light yellow box are (i-k) for Gal-PAMAM and (l-n) for Gal-PETIM at pH 10. Color code: The Gal-dendrimers are shown in orange Licorice format, the buried or internal water molecules are in blue VDW format (c-l, 3 Å), the surface water molecules are in green VDW (d-m, 4 Å), and the bulk water molecules are shown in white QuickSurf format (e-n, 7 Å).

As shown in Fig. 8, the protonated dendrimer shows significantly more water molecules in all the regimes than the non-protonated dendrimers. The internal (or buried) water molecules in protonated dendrimers are notably higher, which suggests its available void volume to accommodate water molecules. The electrostatic repulsion among the branches of the protonated dendrimers causes to solvate more water molecules and subssequently leads to a greater swelling than non-protonated dendrimers. Higher number of water molecules in and around the dendrimers suggest its hydrophilic nature.^80^ It appears that Gal-PAMAM shows more hydrophilic behaviour than Gal-PETIM dendrimer, or conversely, Gal-PETIM dendrimer is more hydrophobic as compared to Gal-PAMAM dendrimer. The observations from the current work are in quantitative agreement with our previous work reported for the PAMAM/PETIM dendrimer.^79,80^ Hydrophobic nature of Gal-PETIM dendrimer can allow better transfection and release of the siRNA to the target cell than Gal-PAMAM dendrimer. These results emphasise that Gal-PETIM dendrimer can be a potential galactose-conjugated dendrimer for the quick release of the genes to the targeted cell.

### PMF Profiles

To quantify the strength of the binding of siRNA and dendrimer at different pH for both the PAMAM and PETIM dendrimer, we have computed the potential of mean force (PMF) using US simulations. The PMF profiles for the complexation between siRNA and Gal-PETIM/Gal-PAMAM (Fig. 9) reveal the influence of pH on binding affinity. The binding free energy (Δ*G*) between siRNA and Gal-PETIM/Gal-PAMAM can be directly calculated from the PMF profile by subtracting the minimum PMF value (representing the bound state) from the unbound state plateau. This difference reflects the work required to separate the siRNA from the dendrimer, quantifying the stability of the complex. The binding free energies for all four different cases are tabulated in the Table. 1.

**Table 1:** Binding free energy between siRNA and Gal-PAMAM and Gal-PETIM dendrimers determined by MD simulation at physiological pH (*i*.*e*. pH 7) and pH 10.

| system | pH | $\Delta G(kcal/mol)$ |
| --- | --- | --- |
| Gal-PETIM | 7 | -86.06 |
|  | 10 | -39.33 |
| Gal-PAMAM | 7 | -178.61 |
|  | 10 | -23.299 |

**Figure 9.**
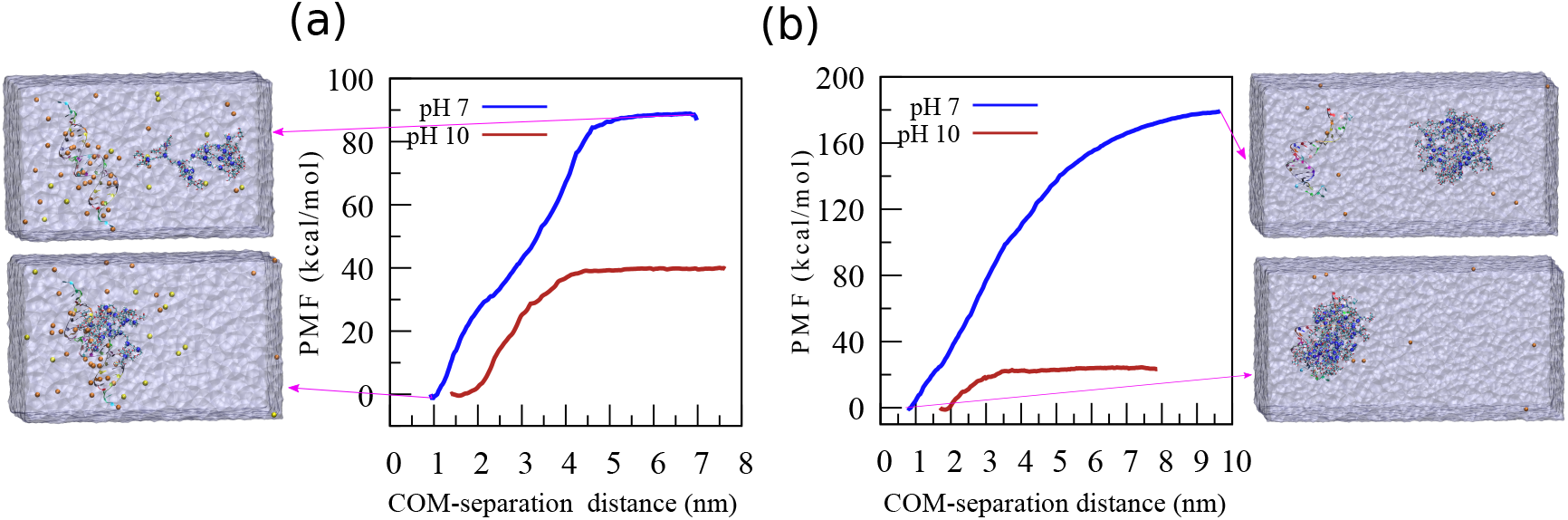
The potential of Mean Force (PMF) profiles for the complexion between siRNA to Galactose-Functionalized (a) G3 PETIM and (b) G3 PAMAM dendrimers along the reaction coordinate. The binding strength for both dendrimers at pH 7 is superior to that at higher pH levels due to the strong electrostatic force of the protonated surface group at physicochemical pH and the negative phosphate group of siRNA. Furthermore, at pH 7, the binding strength is higher due to a larger number of protonated surface groups than PETIM.

The PMF profiles (Fig. 9) revealed greater negative binding free energy for the Gal-PAMAM-siRNA complex than for the Gal-PETIM-siRNA complex at physiological pH, indicating stronger interactions and higher complex stability. This enhanced stability likely arises from stronger electrostatic interactions because of the higher positive charge density of the Gal-PAMAM dendrimer at pH 7. Interestingly, the PMF profiles suggest a reversal of this trend at pH 10, with the Gal-PETIM-siRNA complex exhibiting a more favorable binding free energy.

To gain further insight into the energetic contributions during complexation, we decomposed the classical non-bonded interactions into electrostatic and van der Waals interactions (Fig. 10). As expected, electrostatic interactions dominated the binding process for both complexes at both pH values (Fig. 10a,b). In particular, the Gal-PAMAM-siRNA complex exhibited stronger electrostatic interactions than Gal-PETIM-siRNA at pH 7, consistent with the observed higher binding free energy. In contrast, van der Waals interactions played a lesser role but showed a trend reversal between complexes (Fig. 10b). The Gal-PETIM-siRNA complex displayed greater van der Waals energies at both pH values than Gal-PAMAM-siRNA.

**Figure 10.**
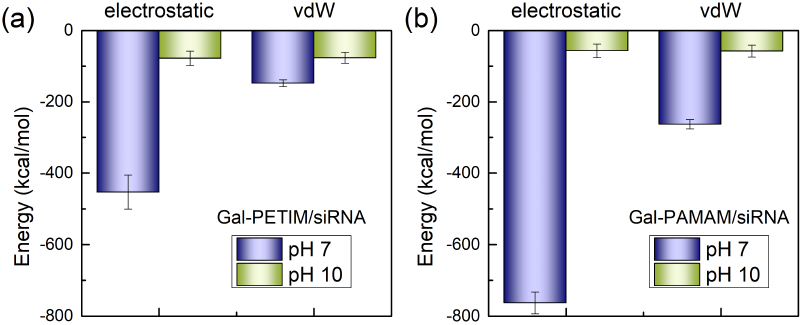
Contributions of electrostatic and van der Waals interactions to the binding of (a) Gal-PETIM/siRNA and (b) Gal-PAMAM/siRNA at pH 7 and pH 10. vdW = Van der Waals.

## Conclusion

In this work, we investigated the complexation of siRNA with galactose-functionalized PAMAM and PETIM dendrimers using allatom MD simulations. We systematically examined the effects of pH on complex formation, and the stability of the resulting complex. Our findings reveal that both the dendrimers readily complex with siRNA, with stronger binding affinity and faster complexation observed at physiological pH (pH 7). This interaction is due to enhanced electrostatic interactions between the protonated dendrimer surface amines and negatively charged siRNA phosphate groups. At higher pH (pH 10), deprotonation of the amines weakens these interactions, leading to less stable complexes. Notably, Gal-PAMAM dendrimers exhibited stronger binding affinity and greater complex stability than Gal-PETIM dendrimers at pH 7, attributed to the higher positive charge density of PAMAM. However, this trend reversed at pH 10, with Gal-PETIM showing more favorable binding free energy compared to Gal-PAMAM. These insights into the pH-dependent behavior and molecular interactions of siRNA-dendrimer complexes provide valuable guidance for the design and optimization of dendrimer-based gene delivery systems for therapeutic applications.

## Author Information

### Authors

**Tarun Maity** - Centre for Condensed Matter Theory, Department of Physics, Indian Institute of Science, Bangalore 560012, India;

**Yogendra Kumar** - Centre for Condensed Matter Theory, Department of Physics, Indian Institute of Science Bangalore, India, 560012;

## Acknowledgments

T.M. acknowledges financial assistance through a MoE scholarship, India. We also acknowledge the computational support provided by the DST-funded TUE-CMS program at IISc.

